# A novel transcriptional signature identifies T-cell infiltration in high-risk paediatric cancer

**DOI:** 10.1101/2022.09.16.508179

**Authors:** Chelsea Mayoh, Andrew J Gifford, Rachael Terry, Loretta MS Lau, Marie Wong, Padmashree Rao, Tyler Shai-Hee, Federica Saletta, Dong-Anh Khuong-Quang, Vicky Qin, Marion Mateos, Deborah Meyran, Katherine E Miller, Aysen Yuksel, Emily VA Mould, Rachael Bowen-James, Dinisha Govender, Akanksha Senapati, Nataliya Zhukova, Natacha Omer, Hetal Dholaria, Frank Alvaro, Heather Tapp, Yonatan Diamond, Luciano Dalla Pozza, Andrew S Moore, Wayne Nicholls, Nicholas G Gottardo, Geoffrey McCowage, Jordan R Hansford, Seong-Lin Khaw, Paul J Wood, Daniel Catchpoole, Catherine E Cottrell, Elaine R Mardis, Glenn M Marshall, Vanessa Tyrrell, Michelle Haber, David S Ziegler, Orazio Vittorio, Joseph A Trapani, Mark J Cowley, Paul J Neeson, Paul G Ekert

## Abstract

Molecular profiling of the tumour immune microenvironment (TIME) has enabled the rational choice of immunotherapies in some adult cancers. In contrast, the TIME of paediatric cancers is relatively unexplored. We speculated that a more refined appreciation of the TIME in childhood cancers, rather than a reliance on commonly used biomarkers such as tumour mutation burden (TMB), neoantigen load and PD-L1 expression, is an essential prerequisite for improved immunotherapies in childhood solid cancers. We combined immunohistochemistry (IHC) and molecular profiling to develop an alternative, expression-based signature associated with CD8^+^ T-cell infiltration of the TIME in high-risk paediatric tumours. Using this novel 15-gene immune signature, Immune Paediatric Signature Score (IPASS), we estimate up to 31% of high-risk cancers harbour infiltrating T-cells. Our data provides new insights into the variable immune-suppressive mechanisms dampening responses in paediatric solid cancers. Effective immune-based interventions in high-risk paediatric cancer will require individualised analysis of the TIME.

## Introduction

The comprehensive genomic analysis of paediatric cancer has provided a wealth of new insights into the distinct molecular nature of childhood cancers. The primary goals of many genomics studies of childhood cancers have focused on the identification of therapeutic options encoded in the genome of cancer cells that would otherwise go unrecognised. Studies such as ZERO Childhood Cancer^1^, INFORM^2^ and the Pediatric Cancer Genome Project^3^ have shown that this can translate into improved patient outcomes. However, the genomic data also includes information about the makeup of the tumour microenvironment, since the RNA of infiltrating immune cells, stromal and vascular cells are also sequenced.

The importance of deciphering the tumour immune microenvironment (TIME) has been driven by the extraordinary impact on the treatment of melanoma, lung adenocarcinoma and head and neck cancers^4^ of agents (typically recombinant antibodies) that inhibit with immune checkpoint molecules such as programmed cell death 1 (PD1), its ligand PD1 ligand 1 (PD-L1) or cytotoxic T lymphocyte antigen 4 (CTLA-4), all of which downregulate T-cell activation^5,6^. TIME and tumour-intrinsic features, such as PD-L1 expression, high tumour mutation burdens (TMB) and high neoantigen load increase the potential for an anti-tumour T-cell response following anti-PD-1 treatment^7^. Evidence also suggests that a more diverse T-cell receptor repertoire is associated with improved response to anti-PD-1^8,9^. However, whilst such biomarkers are applied in some paediatric cancer trials, there is limited evidence to support their validity, and most children with cancer do not respond to Immune Checkpoint Inhibitors (ICIs)^10,11^. This suggests strongly that the microenvironment of childhood cancers is distinct from common adult cancers, and successful immune therapies will require a better understanding of the make-up of the immune microenvironment in childhood cancer types.

The TIME comprises a diverse range of CD45^+^ leukocytes, collectively called tumour infiltrating leukocytes (TILs), which includes CD4^+^ and CD8^+^ T-cells, B-cells, tumour-associated macrophages, dendritic cells, myeloid derived suppressor cells and natural killer cells. The TIME can be classified as ‘immune-inflamed’ with infiltrating T-cells, ‘immune-excluded’ where T-cells are present but confined to the tumour periphery, or ‘immune-desert’ which denotes the total absence of T-cells^12^. Immunohistochemistry is the most established and direct methodology to detect presence of immune cells in a tumour and to assess the relationship between immune and tumour cells. However, RNA sequencing (RNA-seq) is increasingly used to identify immune cell subsets in a tumour and to infer from gene expression profiles the probable nature of the TIME.

Clinical data from immunotherapy trials in adult cancers indicates that T-cell inflamed tumours are more responsive, and this has been a driver to develop multiple different expression-based signatures to detect and characterise TILs^13-17^. Algorithms such as CIBERSORTx (CSX), quanTIseq, and MCP-counter use the expression of key genes to infer the immune cell composition of the TIME from bulk RNA-seq data^18-22^. In contrast to adult cancers, TMB, PD-L1 expression, TIL signatures and deconvolution algorithms have not been systematically applied in paediatric cancers to predict tumour T-cell inflammation. Therefore, a key unmet challenge is identifying the molecular features of paediatric cancers which most accurately characterise the TIME. Two such biomarkers, TMB and PD-L1 expression appear to be less relevant in the paediatric setting than in adult cancers, with the exception of rare hypermutated tumours harbouring a DNA mismatch repair deficiency or polymerase proofreading deficiency. Whilst hypermutated tumours have been shown to respond to immunotherapy^23^, most paediatric tumours have an order of magnitude fewer mutations than in adult cancers^1,24^. PD-L1 expression is reportedly low across paediatric tumour subtypes and the correlation between transcript levels and protein expression is not well established^25,26^, even whilst PD-L1 RNA expression is a trial entry criterion^11^.

We have undertaken a comprehensive characterisation of T-cell infiltration in a diverse spectrum of high-risk paediatric cancers, combining IHC, RNA-seq and whole genome sequencing (WGS). We have cross-referenced the genomic and RNA-seq data with CD8^+^ and CD4^+^ IHC staining on the same tumour specimens to define and validate a novel paediatric specific gene signature that identifies tumours infiltrated by CD8^+^ T-cells. Moreover, we explored the relationship between the gene signature we identified and transcriptional features associated with distinct immune archetypes, providing a unique and detailed insight into the molecular features of the immune landscape across a broad range of paediatric cancers. We also show that PD-L1 RNA expression correlated poorly with PD-L1 protein expression, and that commonly used deconvolution algorithms had only weak correlations with IHC-determined measures of T-cell infiltration. We propose that our novel signature provides a unique and more accurate identification of T-cell infiltrated paediatric cancers.

## Results

### Deconvolution algorithms poorly distinguish individual cell types in high-risk paediatric solid tumours

The ZERO Childhood Cancer Program sequences high-risk paediatric cancers (<30% chance of survival) to identify potential molecularly targeted treatments^27^. The ZERO cohort includes diverse cancer subtypes at various treatment stages -diagnosis, refractory, relapsed or secondary disease (Extended Data Fig. 1). We assigned tumours first into broad disease groups: tumours of the central nervous system (CNS) (N=143), extracranial solid tumours (N=148), or haematological malignancies (HM) (N=56). We further subdivided tumours in these broad categories by tumour subtype (Extended Data Fig. 1). RNA-seq and WGS was performed on 347 samples.

Deconvolution algorithms utilise the expression of key marker genes in bulk RNA-seq data to estimate the relative proportions and types of immune cells present in a sample. We applied CSX, quanTIseq and MCP-counter deconvolution algorithms to identify which tumours might have higher proportions of leukocytes, in particular CD8^+^ T-cells. Estimations of CD8 T-cell abundance were comparable between algorithms (Extended Data Fig. 2a-c), so we subsequently focused on CSX. Unsurprisingly, predictions in haematological malignancies were consistent with the malignancy subtype – myeloid cells predominating in acute myeloid leukaemia, B-cells in B-acute lymphoblastic leukaemia and T-cells in T-cell leukaemia (Fig. 1a-b). All CNS samples had a low total immune cell numbers (median=2.6 cells; range 1.2-7.1). Five extracranial tumours had relatively higher immune cell abundance (>10) than other samples (median=2.7; range 1.1-30.6; Fig. 1a). The predominant immune cell type in solid tumours (CNS and extracranial) was M2 macrophages with lymphocytes making up less than 30%, on average, of the total immune cell populations (Extended Data Fig. 2d-e). However, there were notable exceptions. Twenty-five percent of neuroblastoma (NBL) had predominant monocytes populations, as did some Ewing Sarcomas (EWS), Wilm’s tumours (WT) and medulloblastomas (MB). A malignant peripheral nervous sheath tumour (MPNST), a NBL and an ameloblastic fibrosarcoma (classified as ‘Sarcoma other’) all had activated mast cells comprising greater than sixty percent of the total immune cell population (Extended Data Fig. 2d-e). In 96% of CNS tumours, CD8^+^ T-cells made up less than 10% (median=5%; range 0-25%) of the predicted immune infiltrating cells (Fig. 1c). More extracranial tumours had CD8^+^ T-cells, with 24% of samples having at least 10% (median=5%; range 0-32%) of the immune infiltrating cells predicted to be CD8^+^ T-cells (Fig. 1d). Thus, RNA-seq deconvolution of the immune landscape indicates that T-cell infiltrated paediatric tumours are rare.

**Fig. 1.**
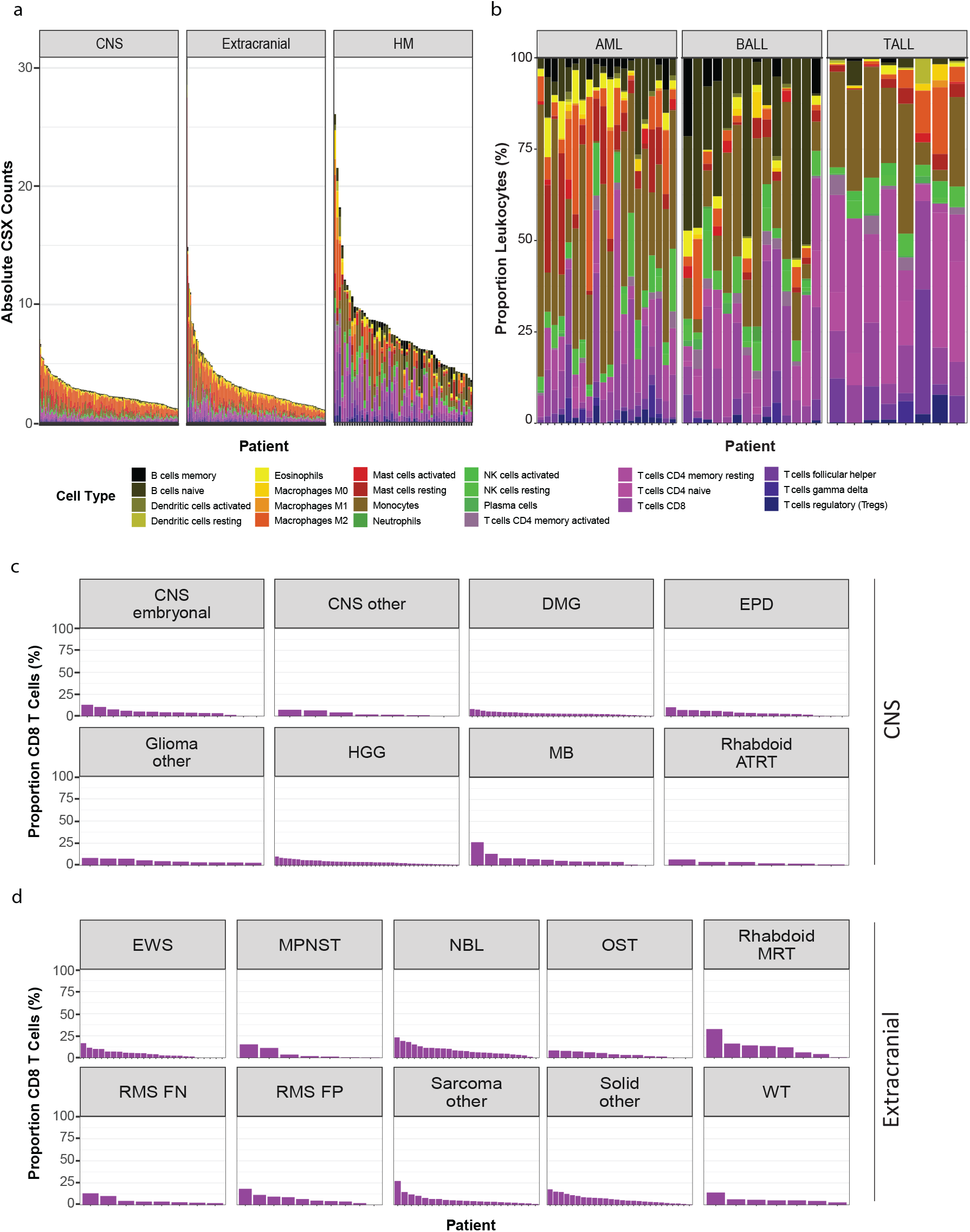
Deconvolution of bulk RNA-sequencing in paediatric cancer. **(a)** Absolute immune cell abundance by CIBERSORTx (CSX) for each patient separated into central nervous system (CNS), extracranial, and haematological malignancies (HM), ordered from highest number of leukocytes to lowest. **(b)** Proportion of all leukocytes (y-axis) for each HM patient (x-axis) separated into acute myeloid leukaemia (AML), B-precursor acute lymphoblastic leukaemia (BALL) and T-cell acute lymphoblastic leukaemia (TALL). **(c** and **d)** Proportion of CD8 T-cells in CSX within each patient classed by CNS **(c)** and extracranial **(d)** tumour subtypes.

### High-risk paediatric cancers are predominantly negative for PD-L1 protein expression

We next explored the relationship between PD-L1 transcript abundance and PD-L1 protein expression in a subset of 59 tumours (20% of cohort) in which we could perform PD-L1 immunohistochemistry. These included both CNS and extracranial tumours. The samples were independently reviewed by an experienced pathology team, blinded to RNA-seq data, and classified as either PD-L1^+^ (≥1 % cells) or PD-L1^−^ (<1% cells) using standard clinical criteria^28^ (Fig. 2a). Only three samples were definitively PD-L1^+^ by IHC, one of which had extremely low levels of PD-L1 mRNA (0.69 TPM; Fig. 2b-c). Of the nine tumours with PD-L1 mRNA expression >3 TPM, only two were PD-L1^+^ by IHC (Fig. 2c). This suggests that mRNA-based thresholds for identifying PD-L1^+^ paediatric tumours may not reliably identify tumours which are PD-L1^+^ by IHC criteria, and PD-L1 TPM based criteria in clinical trials of checkpoint inhibition require further validation.

**Fig. 2.**
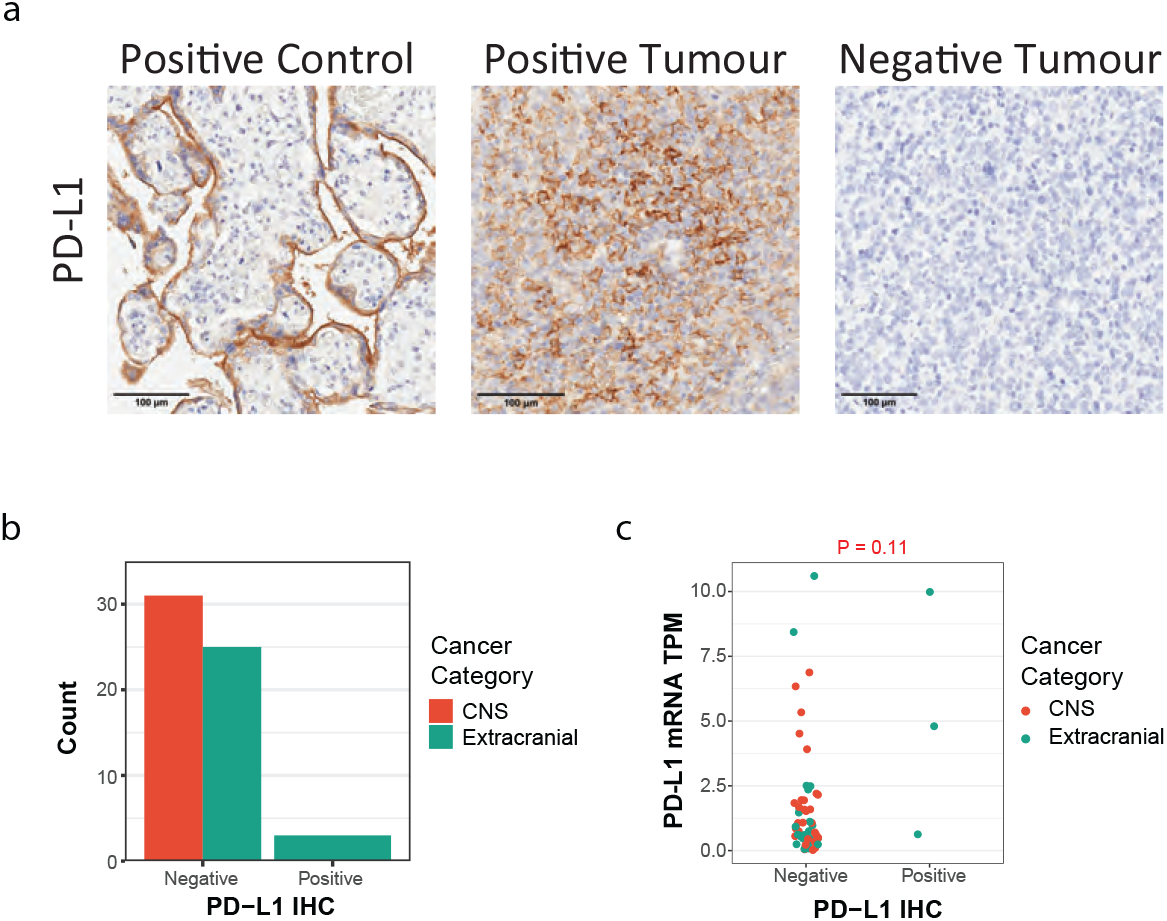
High-risk paediatric cancers are predominantly PD-L1 negative. **(a)** Representative immunohistochemistry (IHC) images for PD-L1 in a control, positive and negative tumour. **(b)** Number of PD-L1 negative and positive samples with cancer category highlighted. **(c)** PD-L1^−^ (<1%) and PD-L1^+^ (≥1%) by IHC compared to PD-L1 expression by RNA-seq (TPM; transcripts per million) in CNS (red) and extracranial tumours (blue).

### Immunohistochemistry identifies immune hot and altered paediatric tumours

We next investigated tumour infiltration by CD45^+^ cells, CD8^+^ T-cells, and CD4^+^ T-cells using RNA-seq deconvolution, and independent analysis of IHC staining for these markers. Seventy-eight samples, representing 27% of the cohort, were analysed (Fig. 3a, Extended Data Fig. 3a-c). We used image analysis to quantitate the number of positive staining cells per mm^2^ (Fig. 3b). In addition, each sample was independently reviewed by a pathologist for CD8^+^ and CD4^+^ infiltration, blinded to the computational or image analysis results. Tumours were classified as either immune inflamed (‘hot’), non-inflamed (‘cold’) or immune-excluded (‘altered’) using published criteria^13^ (Fig. 3c). One sample, without an adjacent haematoxylin and eosin section, was classified as indeterminate. Seventy-six percent (29/38) of CNS tumours were classified as cold, none as hot, and 24% (9/38) were altered. Fewer extracranial tumours were cold (23/39, 59%), 14 were altered (36%) and two hot (5%). Thus 41% classified as either altered or hot (Fig. 3c). We observed a weak positive correlation between the IHC estimates of total leukocytes to the absolute immune cell abundance predicted by CSX (P=0.01, r=0.26; Fig. 3d). However, there was no significant correlation between the computational and pathological estimates of the number of CD8^+^ T-cells (P=0.15, r=0.16) or CD4^+^ T-cells (P=0.8, r=0.03; Fig. 3e-f). This suggested that CSX is not sufficiently sensitive to distinguish individual immune cell types in paediatric samples characterised by low numbers of infiltrating immune cells. The poor correlation between histological and computational predictions of CD8^+^ T-cell abundance was also true of quanTIseq and MCP-counter (Extended Data Fig. 3d-e). The CSX estimate of CD8^+^ T-cell abundance in the two hot samples were the highest estimates in the extracranial cohort, whereas the estimated CD8^+^ T-cell abundance in altered and cold samples were similar (Extended Data Fig. 3f). Contrasting these data with the accurate leukocyte subtype predictions in haematological malignancies (Fig. 1b) suggests that the computational prediction of the immune microenvironment using the tested algorithms depends on immune cell abundance, which in most paediatric tumours are too low to be reliable.

**Fig. 3.**
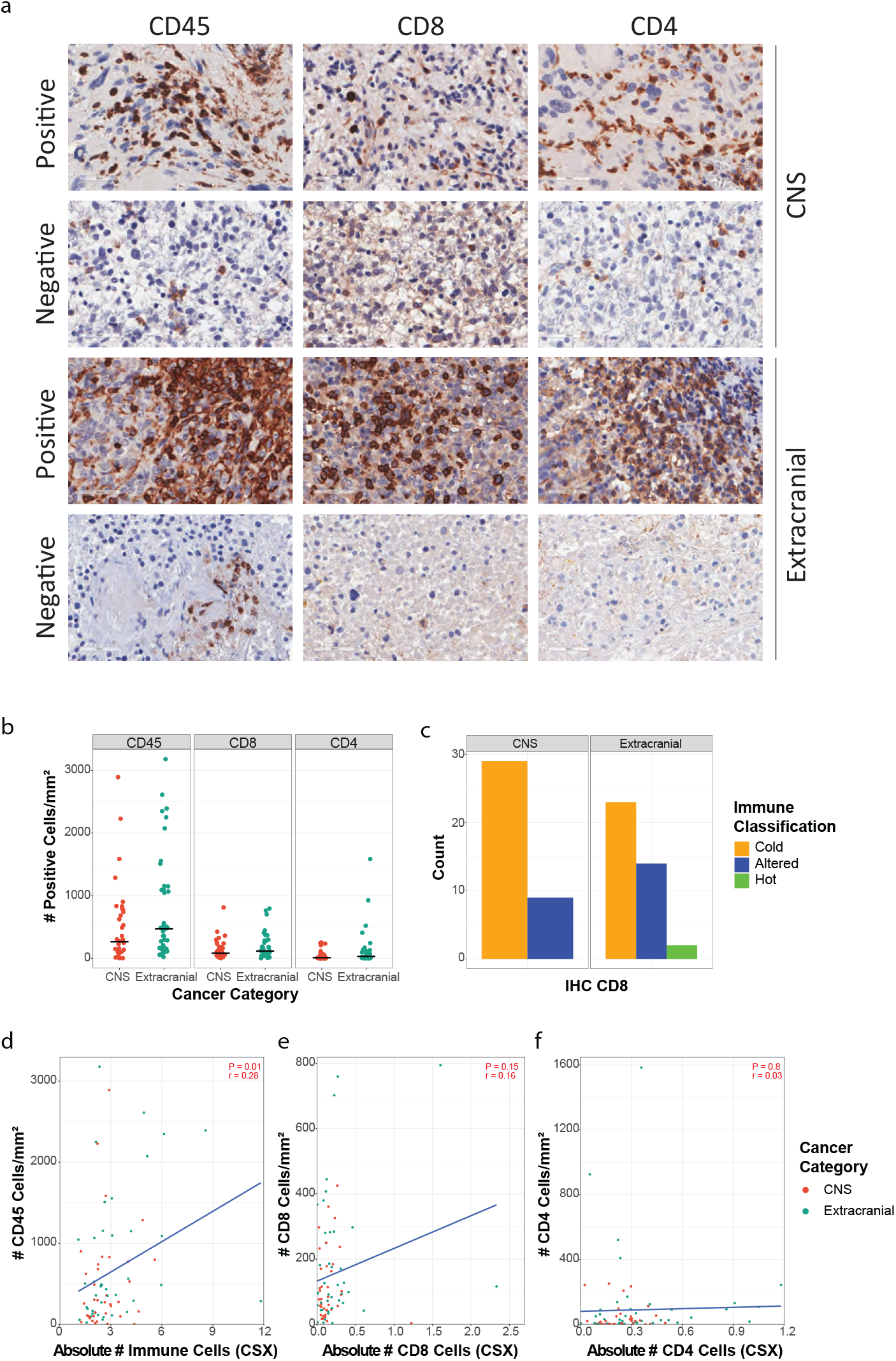
Immunohistochemistry identifies immune-hot and -altered paediatric tumours. **(a)** Representative IHC images for CD45, CD8 and CD4 staining in CNS and extracranial tumours illustrating positive and negative tumours. **(b)** The number (cells/mm^2^) of CD45^+^, CD8^+^, and CD4^+^ cells in CNS and extracranial tumours by IHC. Black horizontal line represents the mean. **(c)** Number of CNS and extracranial samples classified by CD8 IHC as either immune-cold, -altered or -hot. **(d)** Correlation between the absolute number of immune cells by CIBERSORTx (CSX) compared to number/mm^2^ of CD45^+^ cells by IHC. **(e)** Correlation between the absolute number of CD8 T-cells by CSX compared to number/mm^2^ of CD8^+^ T-cells by IHC. **(f)** Correlation between the absolute number of CD4 T-cells by CSX compared to number/mm^2^ of CD4^+^ cells by IHC. Blue line in **(d-f)** is the correlation line of best fit.

Corticosteroids, chemotherapy and radiation therapy have the potential to alter the TIME^29^. To explore this, we looked for correlations between these therapeutic interventions and the number of immune cells per mm^2^ in tumours. No significant difference was observed for CD45^+^, CD8^+^ or CD4^+^ cells in patients who had received chemotherapy or radiation treatment within 42 days of biopsy in IHC data (Extended Data Fig. 4a-c). Further, there was no significant association between corticosteroid administration within 7 days of biopsy and lymphocyte numbers in CNS tumour patients, the population most likely to have received this treatment (Extended Data Fig. 3d-f). This indicates that chemotherapy, radiation and corticosteroid administration are unlikely to be confounding variables altering the analysis of the TIME.

### A novel immune signature predicts T-cell infiltration in high-risk paediatric tumours

We next used the IHC partitioning of tumours as immune hot, altered or immune cold to define a transcriptional signature to predict CD8^+^ T-cell infiltration of high-risk paediatric cancers. For this analysis, we clustered immune hot and altered samples together. We applied machine learning (see methods) to generate a 15-gene signature on the training set (N=34) which was further applied to the test set (N=35) (Fig. 4a). We converted this signature into a score (hereafter referred to as the Immune PAediatric Signature Score (IPASS)) and normalised the IPASS to a range of 1 (most inflamed) to -1 (least inflamed). Using an IPASS of ≥ -0.25 to indicate immune hot/altered and < -0.25 to indicate immune-cold, the IPASS score had a positive predictive value of 78%, a negative predictive value of 92%, sensitivity of 84% and specificity of 88%. In the test set, the resultant signature predicted with 87% accuracy the immune classification of the tumours by IHC. This indicates that the IPASS score can independently classify paediatric tumours as immune hot/altered (“T-cell infiltrated”) or immune cold.

**Fig. 4.**
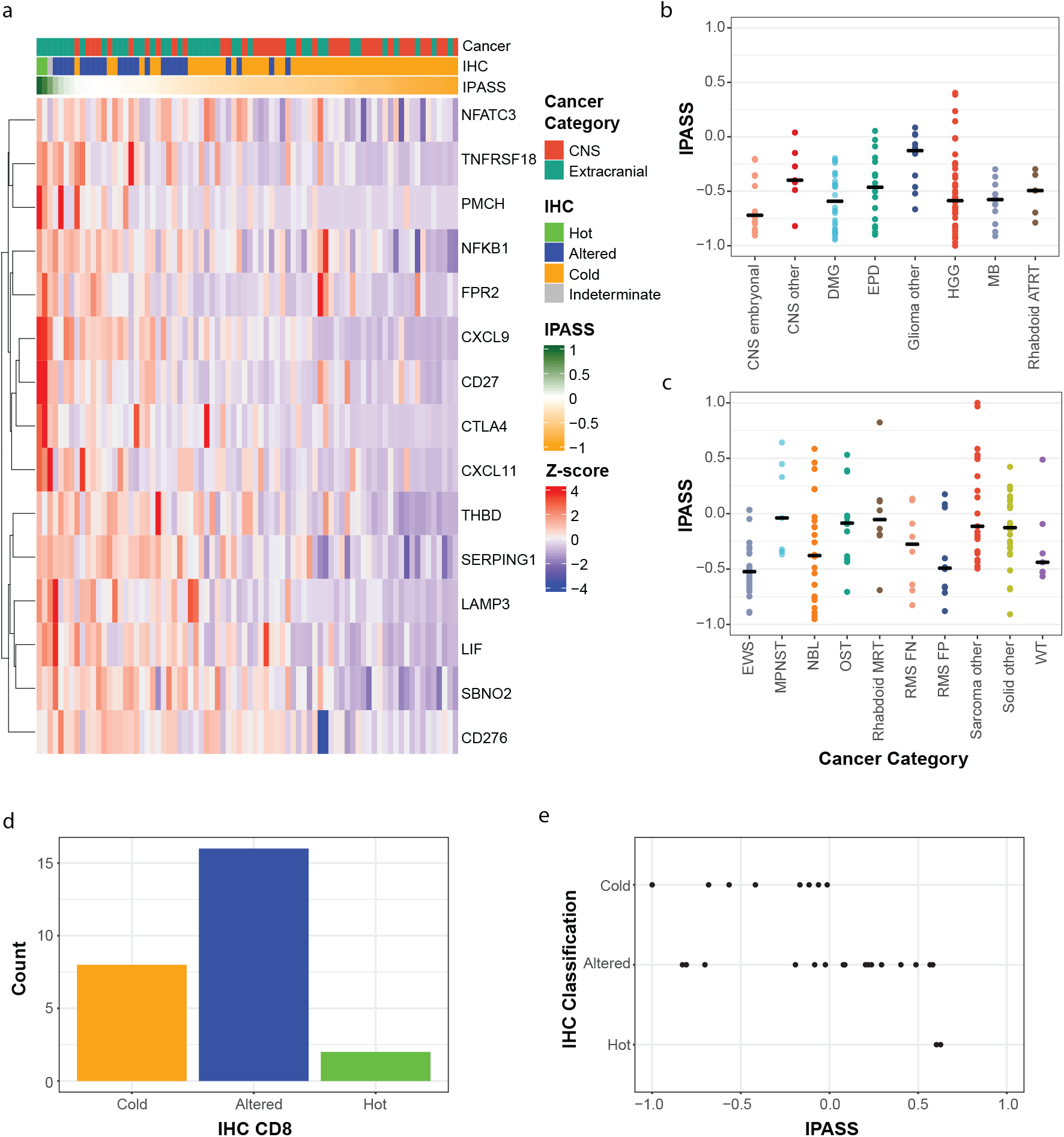
Novel paediatric immune signature predicts T-cell inflamed tumours. **(a)** Heatmap of the novel 15-gene paediatric immune signature (IPASS) in patients with matched IHC (n=78). Top annotation bar represents the cancer category, second annotation is CD8 IHC classification and third annotation bar is the normalised IPASS score measured between 1 (green) and -1 (orange). IPASS distribution across **(b)** CNS and **(c)** extracranial tumours. **(d)** Number of validation samples classified by CD8 IHC as either immune-cold, -altered or -hot. **(e)** IPASS for each validation sample assigned to their IHC CD8 classification (n=26).

This gene signature, constructed to identify immune-hot or immune-altered tumours, incorporates immune markers of inflammation and exclusion (Fig. 4a). Increased expression of *NFATC3, TNFRSF18 (GITR), NFkβ1, CD27 are* associated with T-cell activation. NFATC3 is a transcription factor which initiates production of IL-2 ^30^, *TNFRSF18* and *CD27* are both T-cell costimulatory molecules^31^, NFkβ1 is a key transcription factor generated downstream of T-cell receptor (TCR) signalling^32^ and CTLA-4 is an immune checkpoint expressed following TCR ligation^33,34^. Both *CTLA-4* and *TNFRSF18* are highly expressed on Tregs and increased expression re-enforces the Treg suppressor phenotype^31,35^. *FRP2* (N-formyl peptide receptor 2) is broadly expressed by immune cells and binds several ligands derived from bacterial products leading to initiation of the danger signal response^36^; whereas C1 esterase inhibitor is an inflammatory inhibitor^37^. *CXCL9* and *CXCL11* are key chemokines for trafficking of effector T cells into the tumour and are secreted by macrophages and stromal cells in response to IFN-γ secreted by T-cells^38^. Finally, increased expression of *LAMP3 (DC-LAMP)* and *CD141* (THBD or *BDCA-3*)^39^ are expressed by mature and cross-presenting dendritic cell subsets. The transcriptional repressor *SBNO2* is expressed in macrophages following IL-10/STAT3 signaling and contributes to the anti-inflammatory response^40^. Finally, *B7-H3* expression by tumour cells is immune suppressive, and a current target for immunotherapy strategies^41^.

We applied the IPASS to the remaining samples (n=213) in our cohort. One-hundred and two T-cell infiltrated CNS and extracranial solid tumours were predicted (Fig. 4b-c, Extended Data Fig. 5a-b). Twenty percent (29/143) of CNS tumours had T-cell infiltrated IPASS compared to 49% (73/148) of extracranial tumours (Fig. 4b-c, Extended Data Fig. 5a-b). This is in concordance with the proportions identified by IHC classification (Fig. 3c). T-cell infiltrated tumours were identified across all tumour subtypes except atypical teratoid rhabdoid tumours (ATRT) (Fig. 4b-c). In addition, 21% of high-grade gliomas (HGG) and 15% of diffuse midline glioma (DMG), tumour subtypes sometimes considered non-inflamed^42,43^, had IPASS scores predicting T-cell infiltration (Fig. 4b, Extended Data Fig. 5a). Of the extracranial tumours, relapsed neuroblastoma and malignant peripheral nerve sheath tumours (MPNST) had the highest proportion of immune-infiltrated tumours. Individual examples of epithelioid sarcoma and alveolar soft part sarcoma (both sub-classed as ‘Sarcoma other’) had the highest IPASS (Fig. 4c, Extended Data Fig. 5b). In keeping with our IHC results (Extended Data Fig. 4), IPASS was unaffected by prior treatment or corticosteroid administration (Extended Data Fig. 5a-b).

We tested the validity of the IPASS in an independent dataset (see methods) which underwent bulk RNA-seq and IHC analysis as described for the original cohort. IHC classification in the validation dataset identified 8 cold, 16 altered and 2 hot tumours, which were significantly associated with the IPASS (Fisher’s exact p=0.01; Fig. 4d-e). Extending the IPASS to all samples within the independent dataset that had RNA-seq (N=121) identified twenty-six T-cell infiltrated tumours in an independent dataset (Extended Data Fig. 5c). Taken together, applying the IPASS to ZERO and the validation cohorts suggests that 31% (128/412) of childhood solid tumours are T-cell infiltrated, with the majority falling into the “altered” category, and up to 4% may be true “hot” tumours (Extended Data Fig. 5a-c).

### IPASS correlates with other markers of immune infiltration

T-cell receptor (TCR) clonal diversity within a tumour has been linked to adaptive immune responses as it increased the capacity for T-cells to recognise antigens^44^. For extracranial solid tumours and CNS tumours, we calculated the number of T-cell clones present in each sample from bulk RNA-seq data and tested the correlations with IPASS (Extended Data Fig. 5d-e). The ‘Glioma other’ subgroup (anaplastic pleomorphic xanthoastrocytoma, ganglioglioma and progressive low-grade glioma) had the highest number of TCR clones of the CNS tumours. Osteosarcoma and neuroblastoma had the highest TCR diversity of the extracranial tumours. The number of T-cell clones positively correlated with IPASS (P=1.2e-11, r=0.39; Fig. 5a, Extended Data Fig. 5f).

Elevated tumour-specific neoantigen load is associated with an increased presence of T-cells, particularly in the context of adult tumours with high mutation burdens or childhood cancers arising as a result of germline mutations in mismatch repair genes^23^. This relationship is far less clear in paediatric cancers with much lower mutation burdens. We explored the relationship between the IPASS, TMB and neoantigen burden. There was a wide variation of neoantigen load across tumour types (Extended Data Fig. 6a-b). HGG had the greatest range (5-528), and of the “CNS other” group, 2 choroid plexus carcinomas had over 200 predicted neoantigens (Extended Data Fig. 6a). In extracranial tumours, neoantigen load ranged from 1 to 416, with 4 MPNST samples having >100 predicted neoantigens. High individual neoantigen loads were also observed in an adrenocortical carcinoma and malignant germ cell tumour (both sub-classed as ‘Solid other’) (Extended Data Fig. 6b). As anticipated, a significant correlation was observed between the number of neoantigens and the TMB (P=2.2e-16, r=0.57; Extended Data Fig. 6c). However, neither the TMB nor the number of neoantigens positively correlated with the IPASS (P=0.86, r=-0.01and P=0.96, r=0.0, respectively; Fig. 5a, Extended Data Fig. 6d-e). This suggests that the quantity of mutations and neoantigens is not predictive of T-cell tumour infiltration in paediatric cancers which have mutation burdens within the non-hypermutated range (<5mut/MB). Interestingly, there was significant negative correlation between the IPASS score and estimated tumour purity using WGS data (P=1.6e-14, r=-0.43; Extended Data Fig. 6f). This might be expected if the TIME and other non-tumour cells make up a greater proportion of the sequenced sample. Together, these data establish that neoantigen load and TMB are not indicative of T-cell infiltrated phenotype in most paediatric solid tumours.

**Fig. 5.**
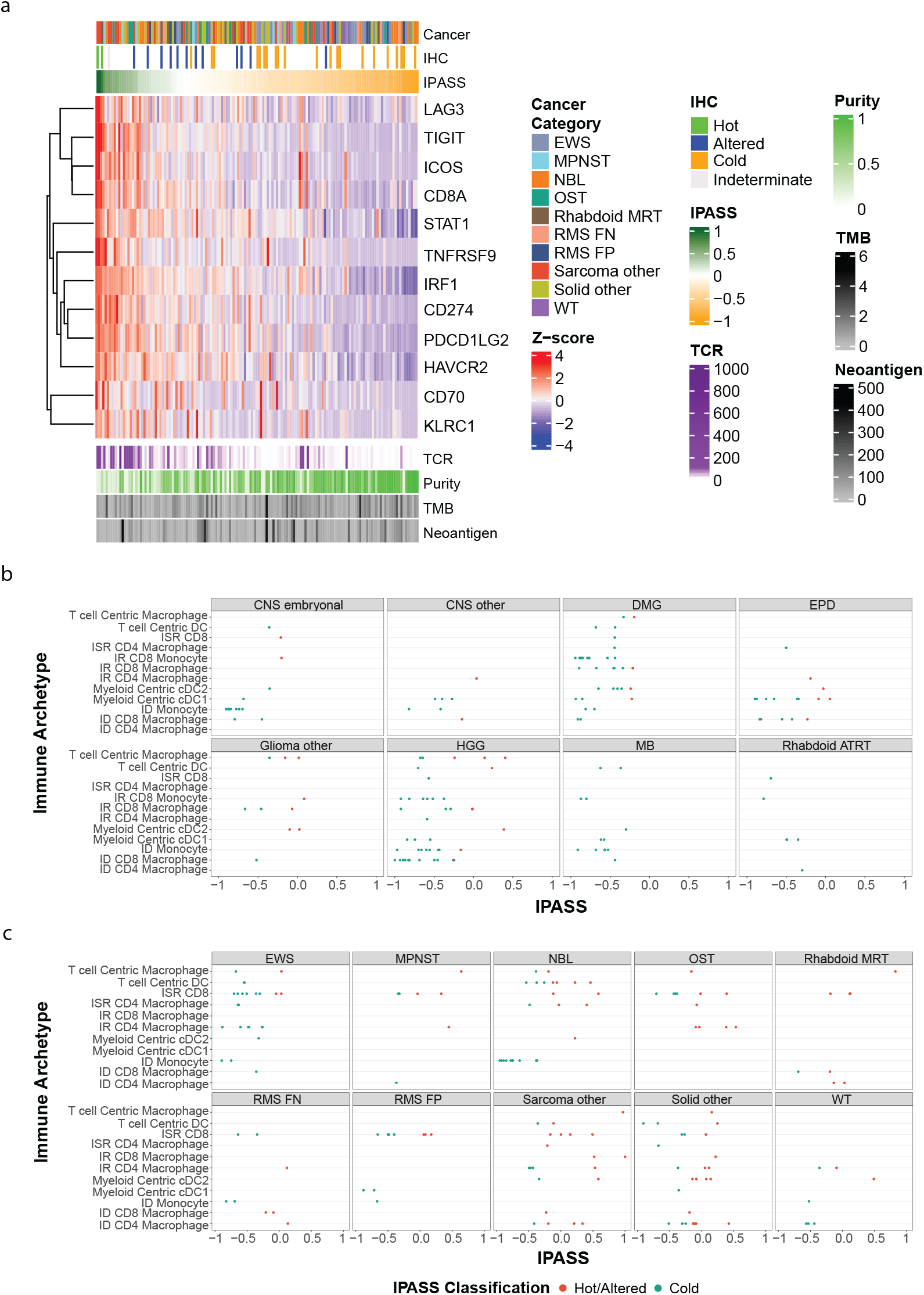
Immune genes and archetypes provide insight into TIME of paediatric tumours. **(a)** Heatmap of 12 immune checkpoint genes associated with IPASS in extracranial tumours (n=148). Top annotation bar represents the cancer category, second annotation is CD8 IHC classification and third annotation bar is the normalised IPASS score measured between 1 (green) and -1 (orange). Below the heatmap, the annotation bars represent the number of T-cell receptor (TCR) clones, tumour purity (percentage of malignant cells), tumour mutation burden (TMB) and neoantigen load. **(b and c)** IPASS score (highlighted by hot/altered (red) or cold (blue)) for each sample assigned to their given dominant immune archetype in **(b)** CNS and **(c)** extracranial tumours. Archetype key: dendritic cells (DC), immune stromal rich (ISR), immune rich (IR), classical DC (cDC), and immune desert (ID).

We next characterized the expression of a selected list of immune checkpoint and regulatory genes (Supplementary Table 1) to identify those most associated with an IPASS >-0.25, in order to understand the potential immune suppression mechanisms regulating T-cell infiltration in high-risk childhood cancers. A subset of 12 genes showed a strong statistical association with high IPASS scores by K-means clustering (Fig. 5a, Extended Data Fig. 6g-h). These include members of the TNF-Receptor super family *CD70* and *TNFRSF9* (CD137, 41BB), Programmed-Death Ligands 1 and 2 (*CD274* and *PDCD1LG2*), Lymphocyte-activation gene 3 (*LAG3*) and T-cell Immunoglobulin genes *TIGIT* and *TIM3* (Fig. 5a). Taken together, increased expression of these genes is a further indication that elevated IPASS is indicative of T-cell activation within these tumours. In CNS tumours these 12 immune genes were weakly associated with higher IPASS scores, in part because fewer CNS samples had scores indicative of T-cell infiltration (Extended Data Fig. 6g).

T-cell tumour infiltration may be indicative of several distinct immune archetypes, both immune rich (IR) and immune desert (ID), within the TIME^45^. We used expression signatures associated with immune archetypes to explore the associations between the IPASS and IR and ID immune archetypes (Fig. 5b-c; Extended Data Fig. 6i-j). Higher IPASS scores were significantly more likely to be associated with IR archetypes (p=0.0026). Most CNS tumours were ID and dominated by archetypes associated with high numbers of monocytes or macrophage (Fig. 5b). There were more IR archetypes in extracranial tumours but no associations of statistical significance with tumour subtypes, indicating the diverse archetypes across our cohort are not constrained by tumour type. Some trends, such as the predominance of the monocyte archetype in ID neuroblastoma, and the lack of any osteosarcomas with an ID archetype of any sort, may prove significant in larger cohorts. However, the association between the higher IPASS scores and IR archetypes indicates that IPASS is identifying immune-infiltrated TIME in most paediatric cancer types. As the detailed nature of the TIME appears to vary at an individual level.

## Discussion

The tumour transcriptome provides high resolution insights into the cellular and molecular basis of individual tumours and the surrounding TIME^1,2^. We set out to characterise the TIME from the sequencing performed in the ZERO Childhood Cancer program, however, most tools that are used to deconvolute the immune signature from bulk RNA-seq data have been developed from adult cancer data sets^19-21^. Thus, we developed a paediatric cancer-specific transcriptional tool, based on a “ground truth” of IHC classification of inflammation status (inflamed, excluded or desert) through integrating these findings with RNA-seq. One important motivation is that many current biomarkers used as surrogates for inflamed tumours, such as TMB, neoantigen load and PD-L1 transcript expression^11,23,46^ have limited applicability in the paediatric setting, where mutation burdens and neoantigen abundance are far lower than in most adult cancers^24^. Thus, whilst we show that neoantigen load correlates with TMB, neither variable correlates with T-cell infiltration. Moreover, if the true HLA affinity threshold is lower than commonly used for neoantigen prediction, then the number of true antigens in paediatric tumours may be even lower than *in silico* predictions^47^. Therefore, beyond the small subset of paediatric patients with hypermutated tumours, an important challenge is to characterise the TIME of paediatric tumours with more typical TMB (<5 muts/Mb). Our data clearly indicate that a proportion of such tumours do harbour infiltrating T-cells. The specific tumour epitopes presented to and recognised by infiltrating T-cells may be more critical for potential T-cell responses than the absolute neoantigen load.

An important relationship we explored is that between PD-L1 gene and protein expression on tumour cells or antigen presenting cells. The importance of this relationship is emphasised PD-L1 gene expression as a criterion on which paediatric patients are selected for trials of anti-PD-L1 therapy^11^. Appropriate selection of patients is critical to trial success, and our data may provide one explanation for the limited anti-tumour activity to ipilimumab, pembrolizumab, nivolumab and atezolizumab seen in paediatric patients^48-51^. In our cohort, PD-L1 protein expression cannot be reliably predicted from RNA-seq data. This is in part because the transcript expression is, in most instances of paediatric cancer, very low (median TPM 0.92) and likely below levels where the relationship between transcript abundance becomes a reliable predictor of protein abundance. Low transcript levels of PD-L1 do not necessarily indicate that a tumour lacks PD-L1 protein. Using PD-L1 transcript counts, fifteen percent of our cohort would be eligible for trial inclusion but only two of these had unequivocal evidence of PD-L1 protein expression. Conversely, TPM criteria would have excluded a patient who was, by IHC, PD-L1^+^. Better informed patient selection might improve the generally disappointing results of anti-PD-1 immunotherapies in paediatrics, as is becoming clearer in the use of these agents in hypermutated tumours^23^ and INI-negative tumours^52^.

The IPASS score is primarily to detect T-cell infiltration of high-risk paediatric cancers. Our data suggests that there are subsets of paediatric tumours, thought previously not to be inflamed, which may in fact harbour a diverse T-cell repertoire. The questions this raises are what antigens are these T-cells responding to and what immunosuppressive pathways characterise individual paediatric cancers? The IPASS gene signature not only identified genes indicative of T-cell activation (from the “hot” samples) but also genes involved in the suppression of T-cell responses (from the “altered” samples), across a broad range of childhood cancers. We have represented the IPASS in a conceptual diagram depicting the tumour immune cellular and signalling network (Fig. 6). Thus, whilst increased expression of *NFATC3, TNFRSF18, NFkβ1, CD27, CTLA4* are indicative of T-cell activation^30-34,53^, *TNFRSF18* and *CTLA4* are also highly expressed on Tregs, which are involved in downregulating antitumour immune responses^31,35^. CD70 is expressed on antigen presenting cells and co-stimulates T-cell via binding to CD27^53^. CD137^54^ is also a costimulatory molecule expressed on activated T-cells within tumours, along with the immune checkpoints LAG3, TIM3 and TIGIT^55^. Whilst PD-L2 is constitutively expressed by antigen presenting cells^56^, PD-L1 is upregulated on tumour cells and antigen presenting cells following IFN-γ stimulation^57^. The expression of *LIF* (Leukaemia Inhibitory Factor) is a novel feature of the IPASS, as LIF has not previously been identified as a prominent feature of the TIME in paediatric cancer. LIF is a ligand for LIF Receptor (LIFR), and a member of the Interleukin-6 cytokine family^58^. LIF has immunosuppressive functions in some tumour contexts, in part by repressing CXCL9 (also part of the IPASS) and CD8^+^ T-cell infiltration of tumours^59^. Establishing the role of LIF in the TIME of childhood cancers potentially opens up the possibility of combined LIF inhibition and anti-PD-L1 immunotherapy.

**Fig. 6.**
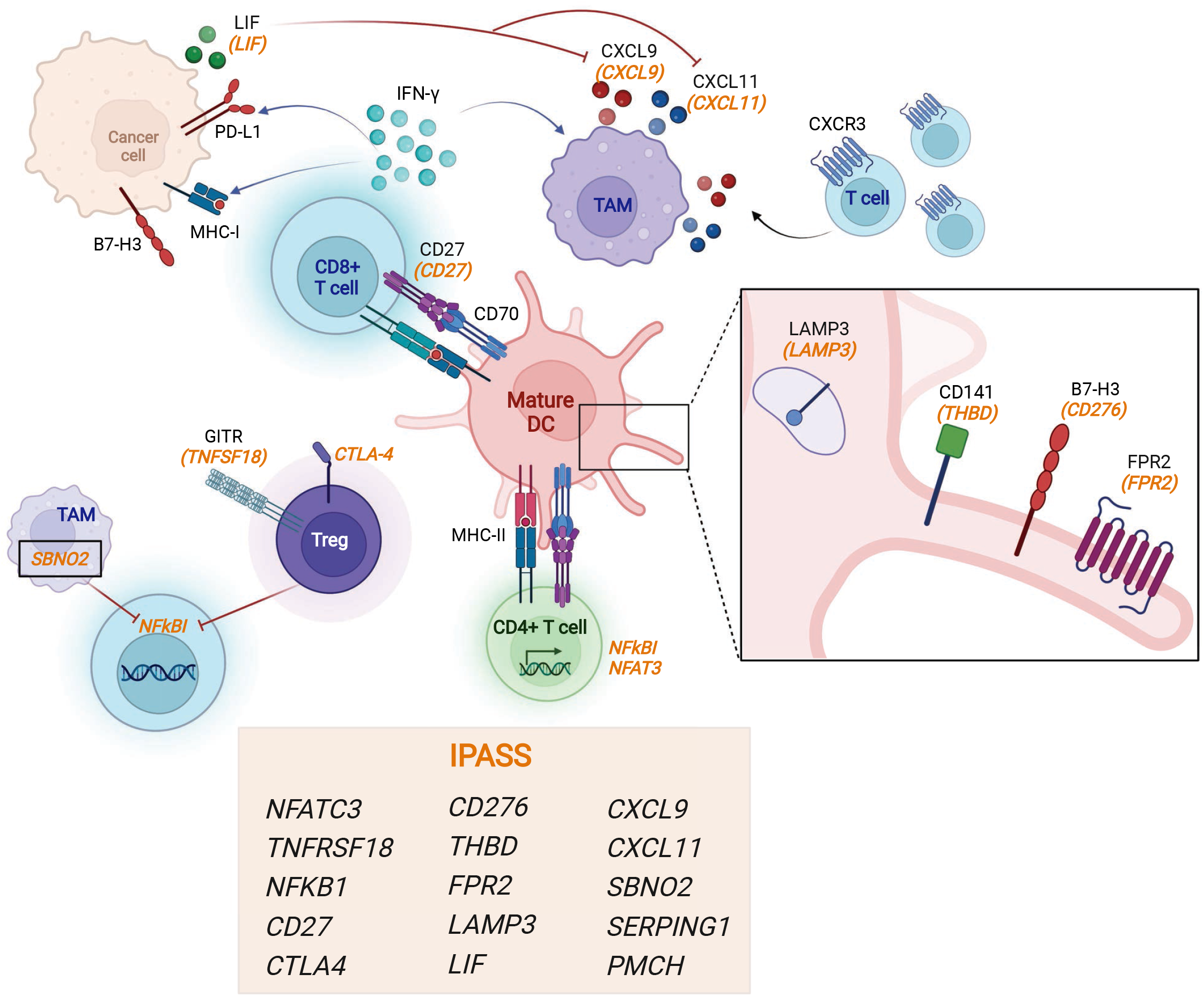
The IPASS gene signature describes a complex immune network within paediatric cancers. A concept figure depicting the potential immune cell interactions which feature the IPASS genes. The IPASS describes interactions which both drive and control anti-tumour immunity. CD8^+^ T-cells (blue) recognize tumour-associated antigen on mature and cross-presenting dendritic cells (pink) *[CD141, LAMP3]* and secrete IFN-g, which induces cancer cell MHC-I and PD-L1 expression. IFN-g response genes [*CXCL9, CXCL11]* are derived from tumour associated macrophages (TAMs, mauve) and are key chemokines for trafficking of CXCR3^+^ effector T-cells into the tumour. Control over this effector T-cell trafficking is mediated by tumour cell secretion of *LIF* which suppresses TAM *CXCL9*. In addition, IL-10/STAT3 signalling in TAMs induces *SBNO2* a transcriptional co-repressor which contributes to the anti-inflammatory response. The IPASS includes genes expressed by activated T-cells *[NFATC3, NFKb1, CD27, CTLA4, GITR]*, in contrast the immune suppressor Tregs (purple) constitutively express *[GITR, CTLA4]*. The functional effect of *B7-H3* is context dependent, *B7-H3* expression on tumour cells is immune suppressive. The transmembrane receptor *FPR2* senses ligands from bacteria products and generates a danger signal. This figure was created in Biorender.

A pan-cancer score like IPASS is necessarily reductive, focused on developing validated and robust ways of characterizing immune-altered and hot tumours across the broad spectrum of paediatric oncology pathologies, but sacrificing some of the detailed individual features of the TIME. Further, the numbers of samples across all tumour subtypes on which we have IHC also limits the resolution of the score. We tried to address this by exploiting other expression-based signatures developed following high-resolution characterizations of the cell type and gene-expression profiles of a broad, adult pan-cancer cohort^45^. This showed, at one level, that our IPASS score is identifying samples with immune-rich signatures, and with a bias towards T-cell infiltration. Further, there are also diverse TIMEs both within tumour subtypes and across the paediatric pan-cancer landscape. Although the numbers of tumours in each subtype limited the capacity to establish definitive links between tumour types and TIME archetypes, it is likely that such patterns will emerge with expanded sequencing of paediatric tumours, particularly if presentation of certain tumour-specific antigens, for example fusion oncogenes, are important in invoking a T-cell response. Combining the IPASS with other approaches such as the immune archetypes, provides a more nuanced and detailed insight into the specific mechanisms operating within individual cancers that distinguish hot and altered tumours from cold tumours, and the complex mechanism of immune evasion. This, we propose, will be the basis of detailed understanding of the immunological features unique to paediatric cancers, and the development of therapeutic approaches that can realise the potential of immunotherapy in solid childhood cancers.

## Methods

### Patients, Samples and Sequencing analysis

All patients and samples were obtained as part of the Australian ZERO Childhood Cancer Precision Medicine Program as described in Wong *et al*. (2020). Specifically, the TARGET pilot program which recruited patients from June 2015 to October 2017 was approved by the Sydney Children’s Hospitals Network Human Research Ethics Committee (LNR/14/SCH/497), and the PRISM clinical trial (NCT03336931) where data was collected from September 2017 until August 2020 was approved by the Hunter New England Human Research Ethics Committee of the Hunter New England Local Health District (reference no. 17/02/015/4.06) and the New South Wales Human Research Ethics Committee (reference no. HREC/17/HNE/29). Informed consent was received for each patient enrolled on the clinical trial. RNA-seq and whole genome sequencing were conducted on all samples from Wong *et al*. (2020) plus an additional 119 samples for a total cohort size of 347 samples. Immune cell-specific IHC was performed on 78 tumour tissue sections from matching patients. WGS and RNA-seq data processing and analysis was as described in Wong *et al*. (2020)^27^. The internal validation cohort of the IPASS on RNA-seq data was performed on an additional 57 patients recruited onto ZERO after the IPASS was developed. An external validation dataset was obtained from The Institute for Genomic Medicine at Nationwide Children’s Hospital (NCH; Columbus, Ohio, USA) for 64 extracranial tumours. Patients at NCH were previously enrolled onto an IRB-approved translational research protocol, which included exome-and RNA-sequencing of frozen tumour as well as banking of additional paraffine embedded tissue blocks. Briefly, RNA-seq libraries were constructed following DNase treatment and ribosomal transcript depletion of total RNA extracts, using the NEBNext® Ultra™ II Directional RNA Library Prep Kit for Illumina (New England Biolabs Inc, Ipswich, Massachusetts, USA), according to the manufacturer’s instructions. Sequencing was performed on either Illumina HiSeq 4000 or NovaSeq 6000 to generate paired 151 base-pair reads, followed by alignment to the GRCh38 reference genome. Transcripts Per Million (TPM) values were generated from aligned paired-end RNA sequence data using Salmon with bootstrapping set to 100^60^.

### Immunohistochemistry

We obtained 78 tumour tissue sections from paraffin embedded blocks of patients with high-risk CNS or extracranial tumours for immunohistochemical analysis (IHC). We received an additional 15 tissue sections from ZERO and 11 from Nationwide Children’s Hospital (NCH) to perform CD8 IHC as a validation cohort. Tissue sections were stained with CD45 (EP322Y, ab40763), CD8 (ab4055), CD4 (EPR6855, ab133616) at a 1:500 dilution and PD-L1 (rabbit monoclonal anit-PD-L1 primary antibody; Cell Signaling Technology, clone: E1L3N, CAT#13684) at a 1:200 dilution on the Leica BOND RX. Human tonsil was used as the positive control for staining, as well as the negative control (secondary antibody only). Tumours were classified as PD-L1 positive if ≥1% of total cells displayed positive membranous staining as described previously^28^. Cytoplastic staining of PD-L1 was also detected, however these cells were not classified as PD-L1 positive tumour cells. CD45, CD8 and CD4 slides were scanned on the Aperio Scanscope XT and analysed using QuPath^61^ software to analyse the number (per mm2) of positive cells in the entire tumour section. Immunohistochemical staining for CD4 and CD8 was further qualitatively assessed by a Paediatric Pathologist (AJG) using a standard light microscope, blinded to the molecularly determined immune status of the tumour. An accompanying H&E-stained slide was available for review for most cases. Tumours were classified as immune “cold”, “altered” or “hot” as previously published^13^. There were few or absent CD4/CD8 positive T-cells in “cold” tumours; more widespread T-cells in “hot” tumours; while “altered” tumours contained either a moderate number of T-cells within the tumour or T-cells at the tumour periphery with absent intratumoural staining.

### Deconvolution Algorithms

CIBERSORTx deconvolution algorithm was performed on the TPM expression matrix derived from RNA-seq data on 347 samples. The ‘impute cell fractions’ job mode was run in both Absolute and Relative mode against the LM22 signature^18^. The analysis was done in B mode for batch correction, quantile normalisation was disabled, and 500 permutations performed. The deconvolution module of quanTIseq was performed on the pre-computed expression matrix (beginning at quanTIseq step 3) with --tumor=TRUE and --method=lsei where deconvolution was performed against 10 immune cell types and the fraction of uncharacterised cells identified. MCP-counter (v1.1) R package was run on the gene expression matrix to identify 8 immune cell populations, endothelial cells and fibroblasts.

### Development of the Novel Paediatric Signature

The IPASS was developed using a Random Forest machine learning approach. Using the IHC classifications we combined the hot and altered together to form an immune-inflamed group. The cohort was randomly assigned into a training set (N=34) and a test set (N=35), with equal proportions of immune-inflamed and -cold in each set. We obtained the expression profile for the training set and filtered for the 766 immune specific genes present in the NanoString immune profiling panel^62^. The classifier was developed using the R packages caret^63^ and randomForest^64^. We then extracted those genes from the classifier (n=15 genes) based on the highest GINI values. We converted this signature into an Immune Paediatric Signature score (IPASS) by calculating the average sum of the log transformed TPM values for the 15-genes in the signature. Using the entire cohort (N=291) we then normalised the IPASS to obtain a score between -1 and 1. In the results we describe the performance of IPASS using a normalised threshold of ≥ -0.25, which corresponds to a non-normalised threshold of ≥ 0.83 to classify samples as T-cell inflamed.

### T-cell receptor sequencing

For identification of T-cell clones we used MiXCR (v3.0.13)^65^ on bulk RNA-seq data using default parameters. Filtering and QC were performed within MiXCR and only clonotypes associated with TCR beta were extracted and used in analysis. The total number of clones, total number of reads and proportions of each clone were assessed for all CNS and extracranial tumours.

### Neoepitope prediction

OptiType (v1.3.3)^66^ was performed on germline paired-end whole genome sequencing fastq files after fishing for HLA reads at 95% identity, taking only the top match with razers3 (v3.5.8) (razers3 -i95 -m 1 -dr 0) to identify HLA-types. Each end was filtered separately before running OptiType with default parameters for paired-end sequences. Somatic variant calls annotated with VEP were combined with the HLA typing for each patient and analysed through pVACseq^67^ using NetMHCcons^68^ for identifying binding of candidate neoantigens for each HLA-type. A mutant peptide was considered for each neoepitope if the mutant peptide had an IC^50^ binding affinity < 500nM, the wild-type peptide had an IC^50^ binding affinity >500nM and was expressed in RNA-seq (TPM>1). Each mutation resulting in a predicted neoepitope was considered as only one neoantigen regardless of the number of predicted neoepitopes. The total number of neoantigens for all CNS and extracranial tumours were assessed.

### Immune gene expression

We sought to explore the expression of a selected list of immune checkpoint and regulatory genes (Extended Data Table 1) and assess the relationship to IPASS. We performed unsupervised hierarchical K-means clustering to identify specific groupings of immune genes where their expression profiles were most correlated with IPASS.

### Immune archetype classification

For deeper immune classification we applied the dominant immune archetypes as defined in Combes *et al*. and followed the gene signature score methods described in the paper^45^. In brief, we downloaded the 12-gene signatures supplied in their supplementary tables and ran the get_score.py function from the papers associated github page (https://github.com/UCSF-DSCOLAB/pan_cancer_immune_archetypes). This calculated a score for each sample for all 12 immune archetypes. The max score for each sample was identified and the tumour was then assigned to this immune archetype.

### Statistics

All statistical analysis and visualisations were performed in R (v3.6.2). All correlation analysis was performed using the Pearson correlation coefficient. The Shapiro-Wilk test was performed to test for normal distribution, F-test for equal variances and the Wilcoxon rank-sum test was used when normally distributed or equal variances were not observed when determining if chemotherapy, radiation or steroid treatment had an effect on T-cell inflammation. A Fisher’s exact test was performed to assess the statistical association between IHC classification and IPASS immune designation of either T-cell inflamed or cold.

## Acknowledgments

We sincerely thank patients and parents for participating in this study. We thank the many clinicians, tumour banks and health professionals for their time acquiring consent for patients and for collection and coordination of samples and associated clinical data at Sydney Children’s Hospital, Randwick; The Children’s Hospital at Westmead; the John Hunter Children’s Hospital; the Queensland Children’s Hospital; the Royal Children’s Hospital Melbourne; the Monash Children’s Hospital; the Adelaide Women & Children’s Hospital; and the Perth Children’s Hospital. Tumour samples and coded data were supplied by the Children’s Cancer Centre Biobank at the Murdoch Children’s Research Institute and The Royal Children’s Hospital (mcri.edu.au/research/projects/childrens-cancer-centre-biobank). Establishment and running of the Children’s Cancer Centre is made possible through generous support by Cancer In Kids @ RCH (www.cika.org.au), The Royal Children’s Hospital Foundation and the Murdoch Children’s Research Institute. We thank ANZCHOG as the trial sponsor and we thank the staff of the Personalised Medicine Theme of the Children’s Cancer Institute for their dedicated work on Zero Childhood Cancer. We thank the Biomedical Imaging Facility (BMIF) at UNSW for access to instruments through which the imaging component of the IHC analysis for this study was carried out. We thank the Australian Federal Government Department of Health, the New South Wales State Government and the Australian Cancer Research Foundation for funding to establish infrastructure to support the Zero Childhood Cancer personalised medicine program. We thank the Kids Cancer Alliance, Cancer Therapeutics Cooperative Research Centre, for supporting the development of a personalised medicine program; Tour de Cure for supporting tumour biobank personnel; The Steven Walter Children’s Cancer Foundation and The Hyundai Help 4 Kids Foundation for supporting G.M.M. and P.G.E.; Samuel Nissen Charitable Foundation for supporting P.G.E.; and the Lions Kids Cancer Genome Project, a joint initiative of Lions International Foundation, the Australian Lions Children’s Cancer Research Foundation (ALCCRF), the Garvan Institute of Medical Research, the Children’s Cancer Institute and the Kids Cancer Centre, Sydney Children’s Hospital. Lions International and ALCCRF provided funding to perform WGS and for key personnel, with thanks to J. Collins for project governance and advocacy. We thank the Cure Brain Cancer Foundation for supporting RNA sequencing of patients with brain tumours; the Kids Cancer Project for supporting molecular profiling and molecular and clinical trial personnel; and the University of New South Wales, W. Peters and the Australian Genomics Health Alliance for providing personnel funding support. The New South Wales Ministry of Health-funded Luminesce Alliance provided funding support for computational personnel and infrastructure. The Medical Research Future Fund, Australian Brain Cancer Mission, the Minderoo Foundation’s Collaborate Against Cancer Initiative and funds raised through the Zero Childhood Cancer Capacity Campaign, a joint initiative of Children’s Cancer Institute and the Sydney Children’s Hospital Foundation, supported the national clinical trial and associated clinical and research personnel. We thank the Cancer Institute of New South Wales and New South Wales Health (fellowship funding for M.J.C.; CINSW Early Career Fellowship 181430 for M.K.M; CINSW Program Grant 2019/TPG2037). We thank the National Health and Medical Research Council (career development fellowship APP1164960 for O.V.). This research was supported by an Australian Government Research Training Program (RTP) Scholarship (for C.M). This research was supported by Tour de Cure and the Australia and

New Zealand Sarcoma Association (funding for R.T.). We thank the 2018 Priority-Driven Collaborative Cancer Research Scheme, co-funded by Cancer Australia and My Room, for personnel and computational support (grant no. 1165556 awarded to M.J.C.). Zero Childhood Cancer is a joint initiative led by Children’s Cancer Institute and the Kids Cancer Centre, Sydney Children’s Hospital, Randwick. K.E.M., C.E.C., and E.R.M. Wish to thank the Nationwide Foundation Pediatric Innovation Fund (Columbus, Ohio, USA).

## Author Contributions

V.T. and E.V.A.M. led and managed the Zero Childhood Cancer Program. M.H., and G.M.M conceived of and designed the Zero Childhood Cancer Project. R.T., V.Q., D.M, D.S.Z, O.V., P.G.E., J.A.T. and P.J.N provided expertise in cellular immunology and immunotherapy. C.M., M.W, R.B.J. and M.J.C. developed and performed bioinformatic methods pertaining to WGS. C.M developed bioinformatic methods pertaining to RNA-seq and performed computational analysis and integration of data. F.A., H.T., L.D.P., A.S.M., W.N., N.G.G., G.M., J.R.H., S.L.K., P.J.W., T.N.T., G.M.M and D.S.Z were clinical leads at recruiting centres and coordinated patient selection. L.M.S.L., D.K., M.K.M., D.G., A.S., N.Z., N.O., H.D., F.A., H.T., and Y.D collected patient treatment information. K.E.M., C.E.C., E.R.M provided additional RNA and patient slides for external validation cohort. A.J.G. provides pathology expertise. A.J.G., F.S., and T.S. analysed IHC data. R.T. and P.R optimised and stained for CD45, CD4 and CD8. A.Y. and D.C optimised and stained for PD-L1. C.M., M.J.C., J.A.T., P.J.N., and P.G.E. conceived, designed and wrote the manuscript with comments and contributions from all authors.

---

**Extended Data Fig. 1. Cohort Overview**

Representation of the cohort consisting of 347 high-risk paediatric cancers who underwent RNA-seq. The cohort is highlighted by the frequency of samples, from the inner most ring to the outer ring by: CNS tumours (CNS), extracranial tumours and haematological malignancies (HM), further divided into histologies, stage of disease (diagnosis, refractory disease, relapse and secondary cancer), and immunohistochemistry (IHC) performed for samples highlighted in grey. Cancer histology key: acute myeloid leukemia (AML), B-precursor acute lymphoblastic leukemia (BALL), Juvenile myelomonocytic leukaemia (JMML), T-cell acute lymphoblastic leukemia (TALL), diffuse midline glioma (DMG), ependymoma (EPD), high grade glioma (HGG), medulloblastoma (MB), atypical teratoid rhabdoid tumour (ATRT), Ewing’s sarcoma (EWS), malignant peripheral nerve sheath tumour (MPNST), neuroblastoma (NBL), osteosarcoma (OST), malignant rhabdoid tumour (MRT), rhabdomyosarcoma fusion negative (RMS FN), rhabdomyosarcoma fusion positive (RMS FP), and Wilms tumour (WT).

**Extended Data Fig. 2. Deconvolution algorithms exhibit high concordance and an abundance of M2 macrophages in paediatric cancer**

Correlation of total CD8 T-cells between **(a)** CIBERSORTx (CSX) and quanTIseq, **(b)** CSX and MCP-counter (MCP) and **(c)** quanTIseq and MCP. Blue line is the correlation line of best fit. Proportion of all leukocytes (y-axis) for each patient (x-axis) separated by diagnosis in **(d)** CNS and **(e)** extracranial tumours.

**Extended Data Fig. 3. Immunohistochemistry identifies paediatric patients with T-cell inflamed tumours**

Representative IHC images of human tonsil, CNS tumours and extracranial tumours expressing high and low numbers of **(a)** CD45^+^ cells, **(b)** CD8^+^ T-cells and **(c)** CD4^+^ T-cells. Correlation between number/mm^2^ of CD8^+^ T-cells by IHC compared to **(d)** total percent of CD8 T-cells by quanTIseq and **(e)** CD8 score by MCP-counter (MCP). **(f)** Absolute CD8 T-cells in CSX separated by IHC classification in extracranial tumours.

**Extended Data Fig. 4. Prior treatment and steroid administration do not significantly affect T-cell inflammation**

The number/mm^2^ of **(a)** CD45^+^, **(b)** CD8^+^ and **(c)** CD4^+^ cells in CNS (red) and extracranial (blue) tumours identified by IHC separated into yes or no having had received chemotherapy or radiation treatment within 42 days (n=76). The number/mm^2^ of **(d)** CD45^+^, **(e)** CD8^+^ and **(f)** CD4^+^ cells by IHC in CNS (red) tumours by IHC separated into yes or no having had received corticosteroid treatment within 7 days (n=37). Horizontal line is the median percent of cells and the ends of the box represent the upper and lower quartiles.

**Extended Data Fig. 5. Distribution of IPASS and T-cell clones are heterogeneous across histologies**

Heatmap of the IPASS in **(a)** CNS tumours (N=143) and (**b**) extracranial tumours (N=148). Top annotation bar represents the cancer category, and second annotation bar represents treatment (within 42 days) (Dark grey; Yes, Grey; unknown, Light grey; No). The third annotation bar in a represents steroid administration (within 7 days) (Dark grey; Yes, Grey; unknown, Light grey; No, White; not applicable (as not a CNS tumour)). The fourth annotation bar in a and third in b is CD8 IHC classification (white are samples with no IHC data). The bottom annotation bar is the normalised IPASS score measured between 1 (green) and -1 (orange). **(c)** Heatmap of the IPASS extended to all samples in the validation cohort that had RNA-seq data from NCH (n=64) and ZERO (n=57). Top annotation bar represents the cancer category, second annotation is CD8 IHC classification (white are samples with no IHC data), and bottom annotation bar is the normalised IPASS score measured between 1 (green) and -1 (orange). Number of T-cell clones across **(d)** CNS and **(e)** extracranial tumours. Horizontal line is the median number of T-cell clones and the ends of the box represent the upper and lower quartiles. **(f)** Correlation between the IPASS and total number of T-cell clones (n=291). Blue line is correlation line of best fit.

**Extended Data Fig. 6. IPASS correlations with additional markers of immune infiltration**

Number of neoantigens distributed across **(a)** CNS and **(b)** extracranial tumours. The red line represents the median number of neoantigens. Correlation between **(c)** total number of neoantigens and mutations/Mb (coding missense SNPs), **(d)** IPASS and mutations/Mb (coding missense SNPs), **(e)** IPASS and total number of neoantigens, and **(f)** IPASS and tumour purity (n=291). Blue line is correlation line of best fit. Heatmap of all immune checkpoint genes examined in **(g)** CNS tumours (n=143) and **(h)** extracranial tumours (n=148). Top annotation bar represents the cancer category, second annotation is CD8 IHC classification (white are samples with no IHC data) and third annotation bar is the IPASS score measured between 1 (green) and -1 (orange). Bottom annotation bar from top to bottom represents the number of T-cell receptor (TCR) clones, tumour purity (percentage of malignant cells), tumour mutation burden (TMB) and neoantigen load. Proportion of tumours broken into diagnosis assigned to their given dominant immune archetype in **(i)** CNS and **(j)** extracranial tumours. Archetype key: dendritic cells (DC), immune stromal rich (ISR), immune rich (IR), classical DC (cDC), and immune desert (ID).

**Supplementary Table 1. Immune checkpoint and regulatory genes**

