## Supplementary figures and images for "A novel transcriptional signature identifies T-cell infiltration in high-risk paediatric cancer"

### Supplementary Figures

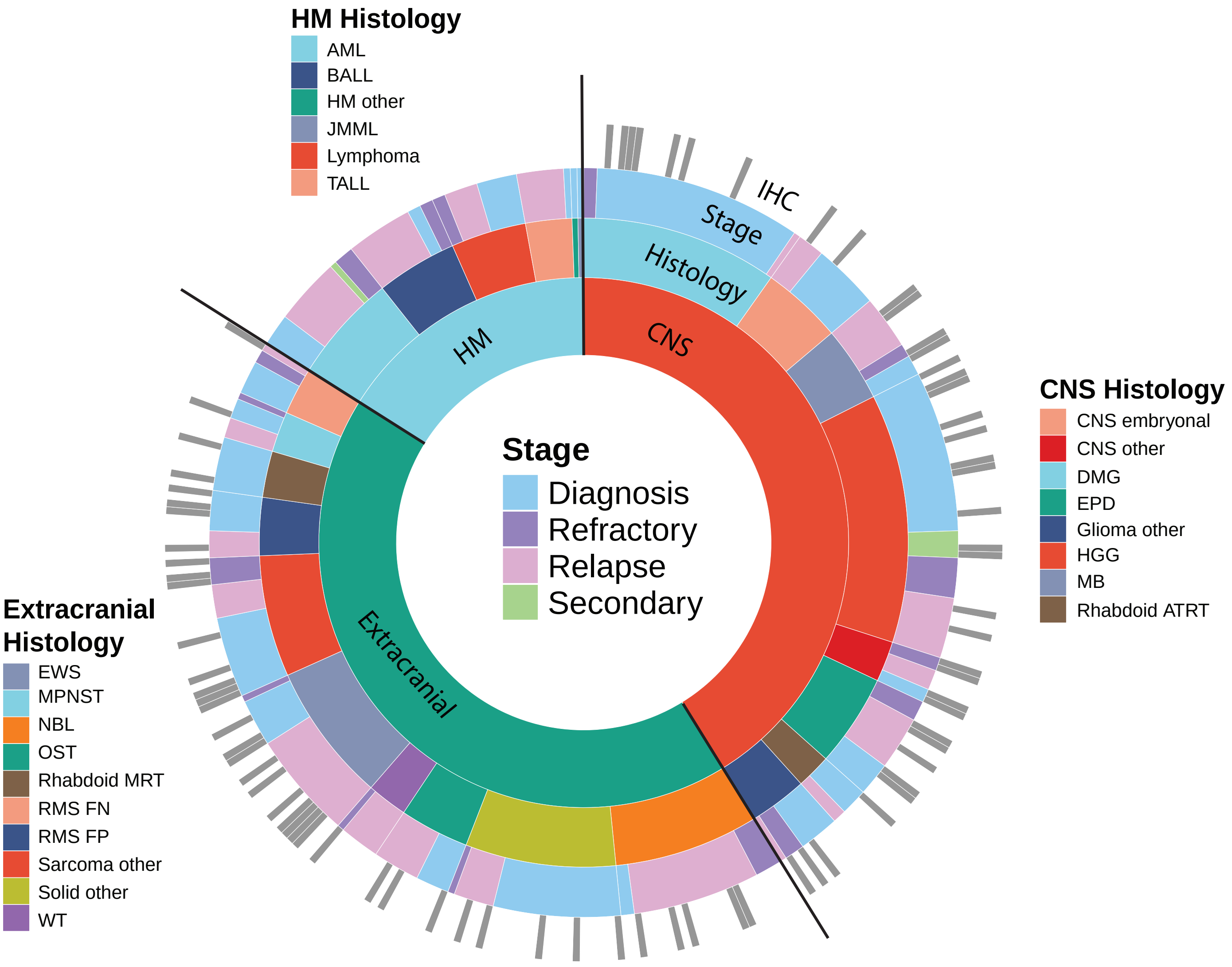

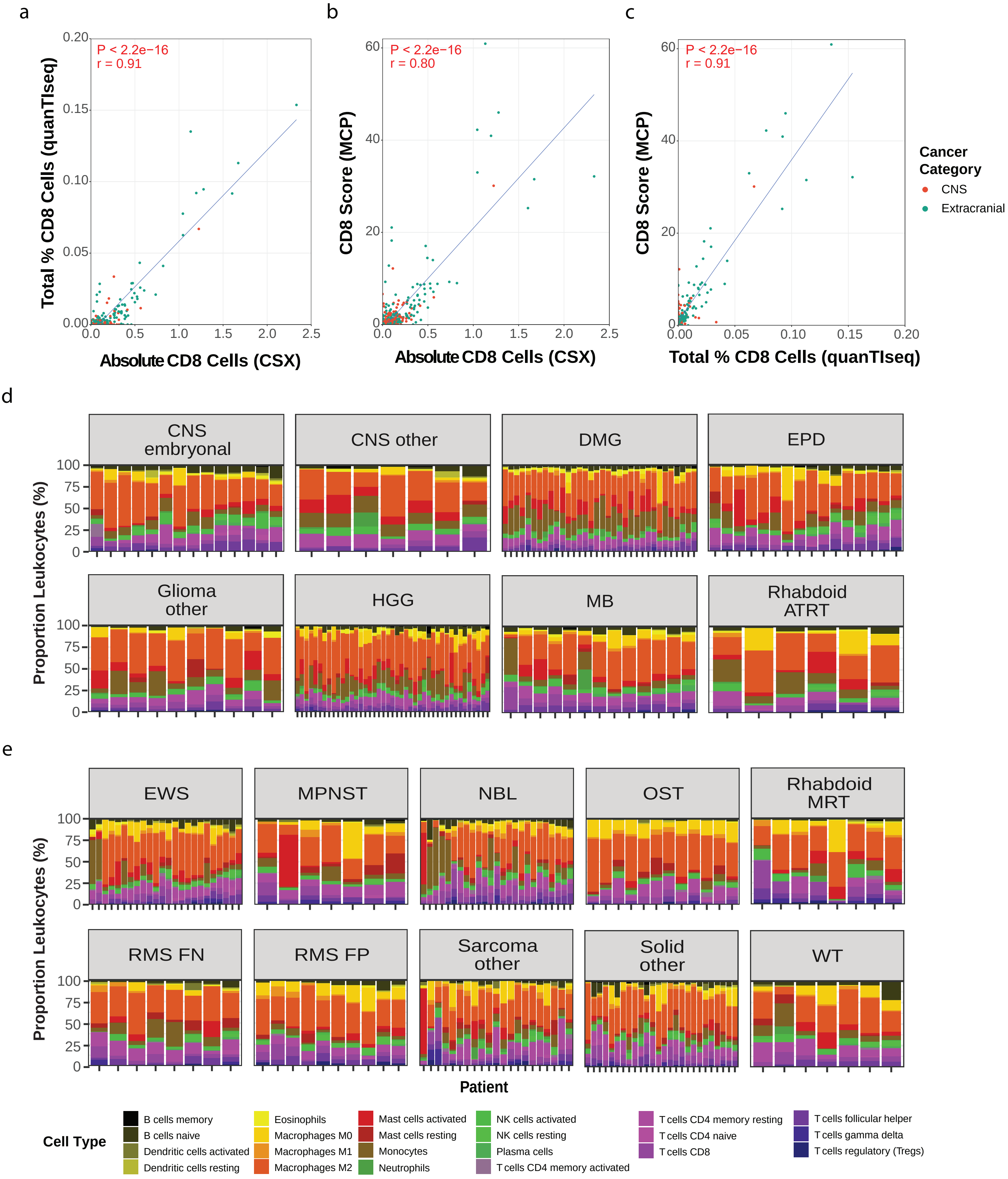


Extended Data Fig. 2

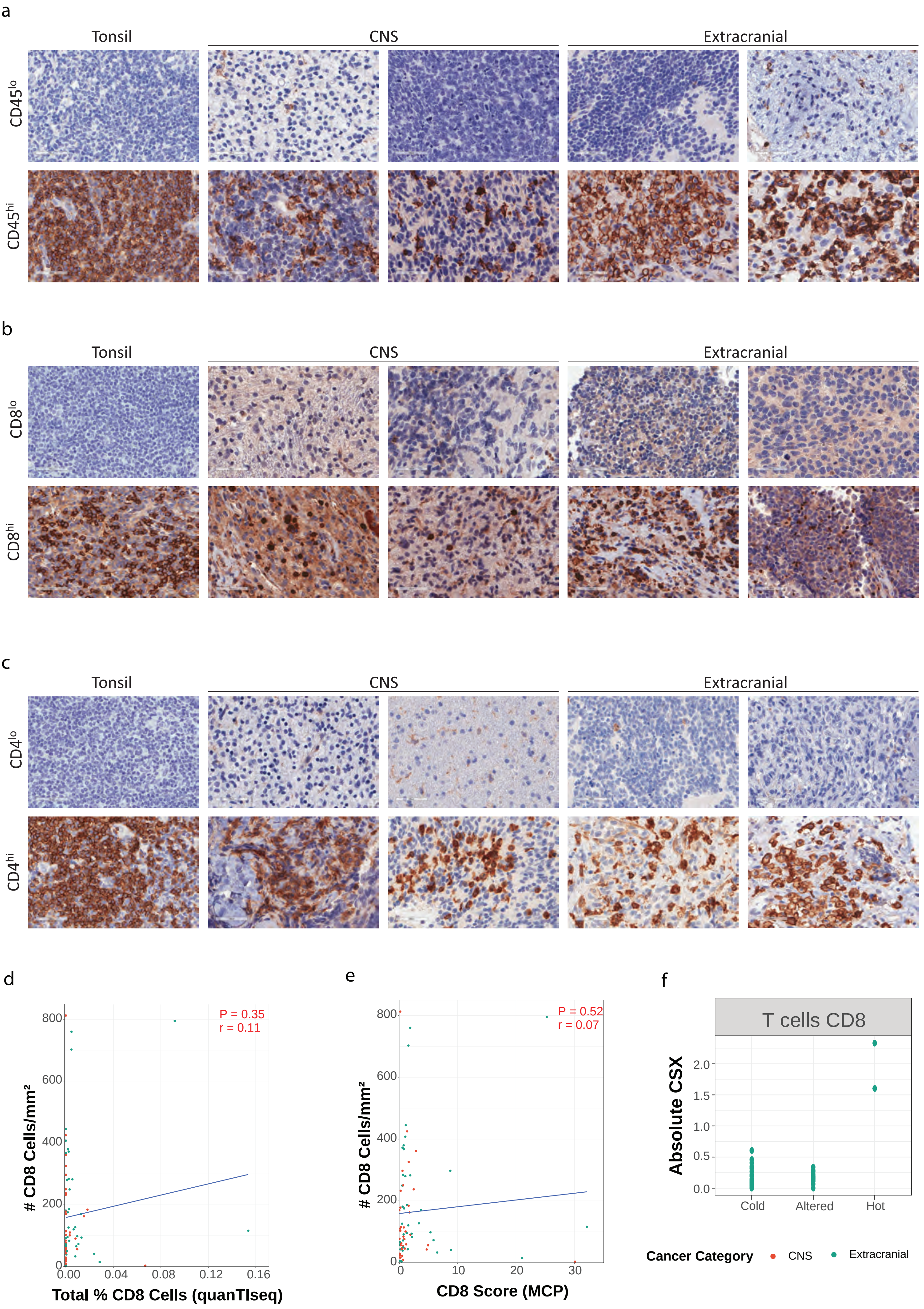


Extended Data Fig. 3

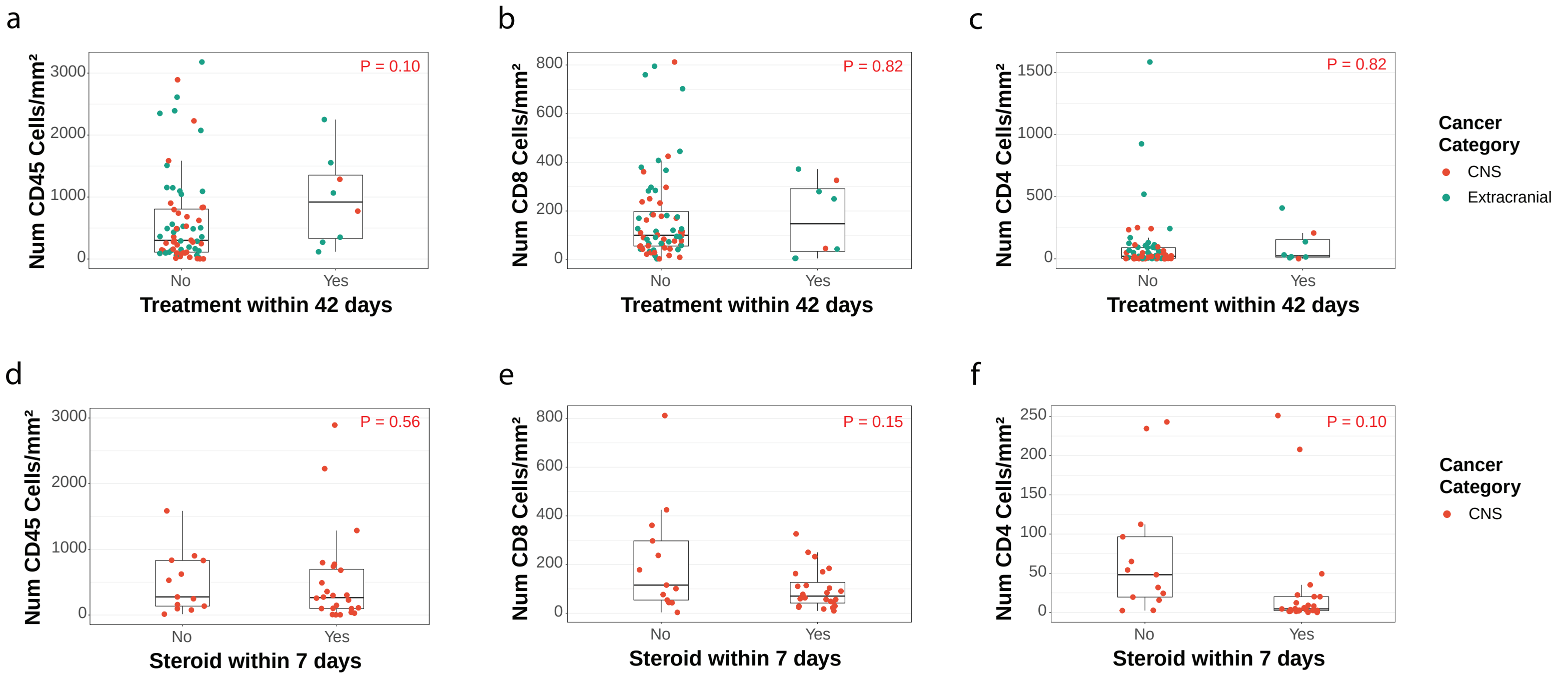


Extended Data Fig. 4

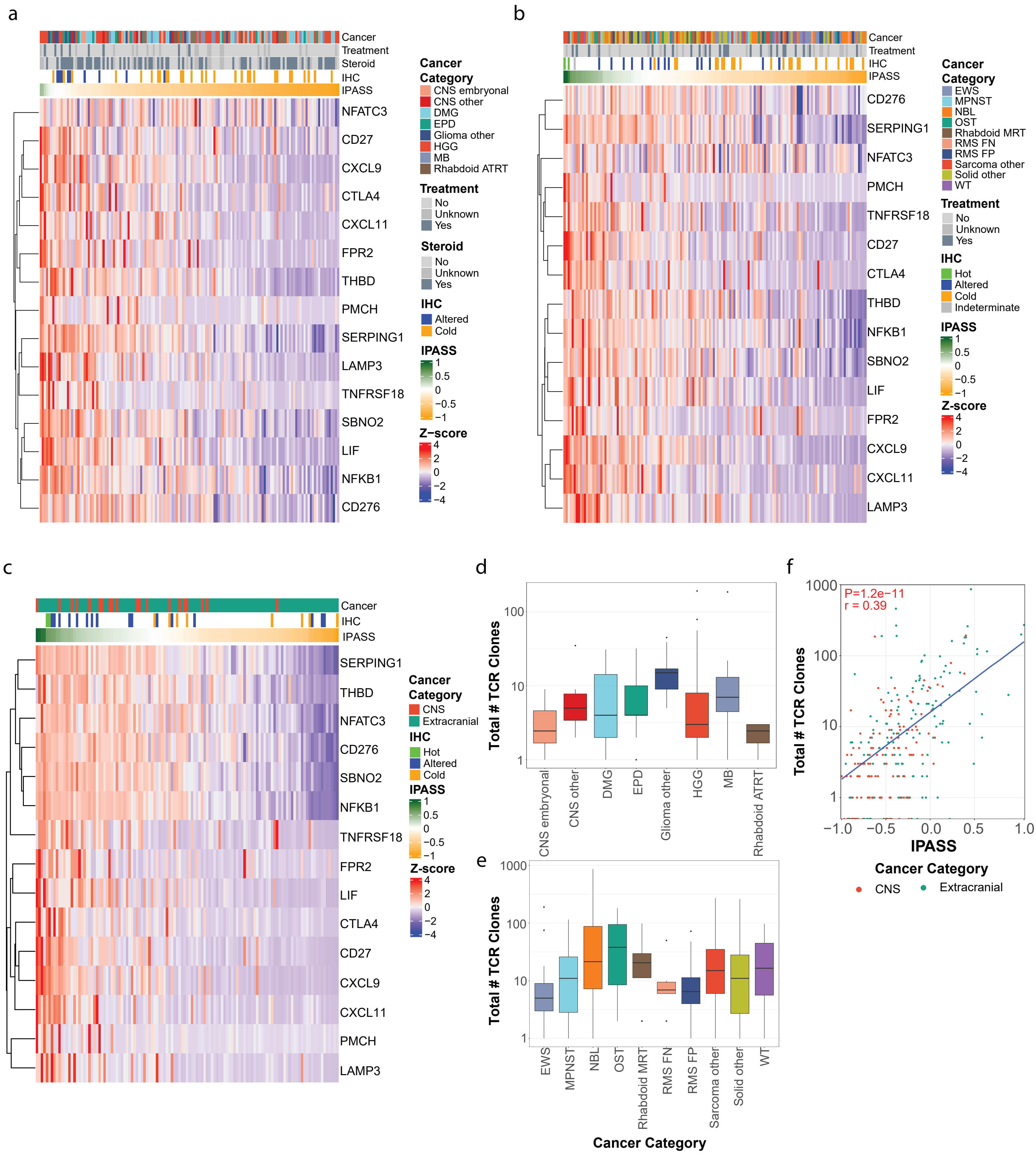


Extended Data Fig. 5

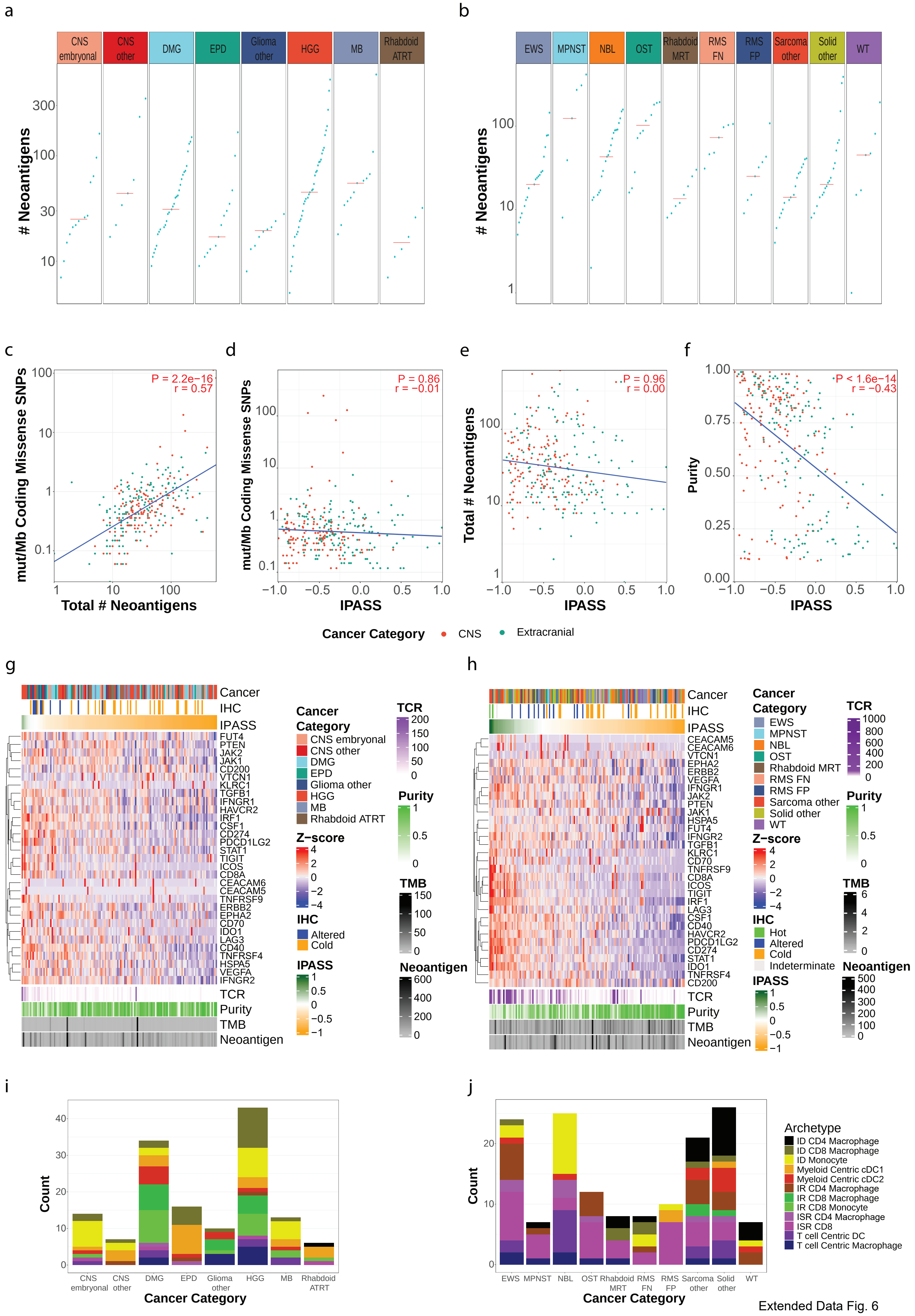


Extended Data Fig. 6
